# Possible link between higher transmissibility of B.1.617 and B.1.1.7 variants of SARS-CoV-2 and increased structural stability of its spike protein and hACE2 affinity

**DOI:** 10.1101/2021.04.29.441933

**Authors:** Vipul Kumar, Jasdeep Singh, Seyed E. Hasnain, Durai Sundar

## Abstract

The Severe Acute syndrome corona Virus 2 (SARS-CoV-2) outbreak in December 2019 has caused a global pandemic. The rapid mutation rate in the virus has caused alarming situations worldwide and is being attributed to the false negativity in RT-PCR tests, which also might lead to inefficacy of the available drugs. It has also increased the chances of reinfection and immune escape. We have performed Molecular Dynamic simulations of three different Spike-ACE2 complexes, namely Wildtype (WT), B.1.1.7 variant (N501Y Spike mutant) and B.1.617 variant (L452R, E484Q Spike mutant) and compared their dynamics, binding energy and molecular interactions. Our result shows that mutation has caused the increase in the binding energy between the Spike and hACE2. In the case of B.1.617 variant, the mutations at L452R and E484Q increased the stability and intra-chain interactions in the Spike protein, which may change the interaction ability of human antibodies to this Spike variant. Further, we found that the B.1.1.7 variant had increased hydrogen interaction with LYS353 of hACE2 and more binding affinity in comparison to WT. The current study provides the biophysical basis for understanding the molecular mechanism and rationale behind the increase in the transmissivity and infectivity of the mutants compared to wild-type SARS-CoV-2.

## Introduction

The SARS-CoV-2 (Severe Acute Respiratory Syndrome-Coronavirus detected in December 2019 in the Wuhan province of China has caused the COVID-19 pandemic. As of April 22, 2021, there are more than 143,184,614 confirmed cases, and 3,047,322 people have lost their lives (https://covid19.who.int/). The SARS-CoV-2 belongs to the family of beta corona virus, the same class of viruses that created the pandemic in the past such as SARS-CoV and MERS ^1, 2^. SARS-CoV-2 possesses a large single-stranded RNA as genetic material and has four main structural components, namely, Envelope protein, Spike protein, Membrane protein and Nucleocapsid ^3, 4^. The main structural element that enables this virus to attach to the host receptor is the spike glycoprotein, and it also gives the crown-like appearance to the virus, hence named as Coronavirus ^5–7^. The Spike glycoprotein of SARS-CoV-2 attaches to the human Angiotensin Converting Enzyme (hACE2) receptor and then activated by another human enzyme, Transmembrane Protease Serine (TMPRSS2), to entry inside the host cells ^7, 8^. Since Spike is the primary target receptor for the entry and the main virulence factor of the virus, various therapeutic drugs and vaccines are being made and tested against it ^9^. Although multiple medications such as remdesivir or hydroxychloroquine lopinavir and ritonavir have been recommended by the World Health Organization (WHO) against COVID-19, their efficacy is still the topic of debate ^10–12^. Similarly, WHO has issued an emergency use listing for certain vaccines such as BNT162B2 from Pfizer, AstraZeneca/Oxford COVID-19 vaccine, manufactured by the Serum Institute of India and SKBio and Ad26.COV2.S, developed by Janssen (Johnson & Johnson) (https://www.who.int/covid-19/vaccines). However, the SARS-CoV-2 cases are still increasing at an alarming rate all over the globe, and the primary rationale behind it is the rapid accumulation of mutations in the SARS-CoV-2.

In the last few months, variants of the SARS-CoV-2 have been reported. Some of them are the Variant of Concern (VOC), which have increased the infectivity or have the potential of immune scape. Almost all the VOCs reported till now have the mutations in the Spike glycoprotein of the virus, which has increased the binding affinity of the virus to hACE2 or has acquired immune scape potential ^13, 14^. The Lineage B.1.1.7 or 20I/501Y.V1 was detected in the United Kingdom in October 2020. This variant has increased the transmissibility by 40-80%, which has been partially correlated with N501Y mutation in Receptor Binding Domain (RBD) of Spike protein ^15^ (**Figure 1A**). In December 2020, B.1.351 was detected in the South African population, which could infect more younger people and had three primary mutations in the RBD of Spike protein, namely, N501Y, K417N and E484K ^16^. Similarly, the lineage P.1 detected in January 2021 in the Brazilian population had three mutations of concern in Spike RBD, namely, N501Y, K417T and E484K ^14, 17^. In our previous study, we had reported that N501Y mutation could enhance the ACE2 affinity and possibly confer resistance towards the antibodies ^14^. Our results also indicated the reinfection potential of P1 and N501Y.V2 variants. In India, lineage B.1.617 (double mutant) and B.1.618 (triple mutant) has been recently reported, which has caused a rapid increase in the COVID-19 cases in the country. In B.1.617 (double mutant), apart from mutations in other proteins, it has three primary mutations in spike glycoprotein, namely, P681R, E484Q, and L452R – the latter two mutations in the RBD region are the Mutation of Concern ^18^. (**Figure 1B**).

**Figure 1.**
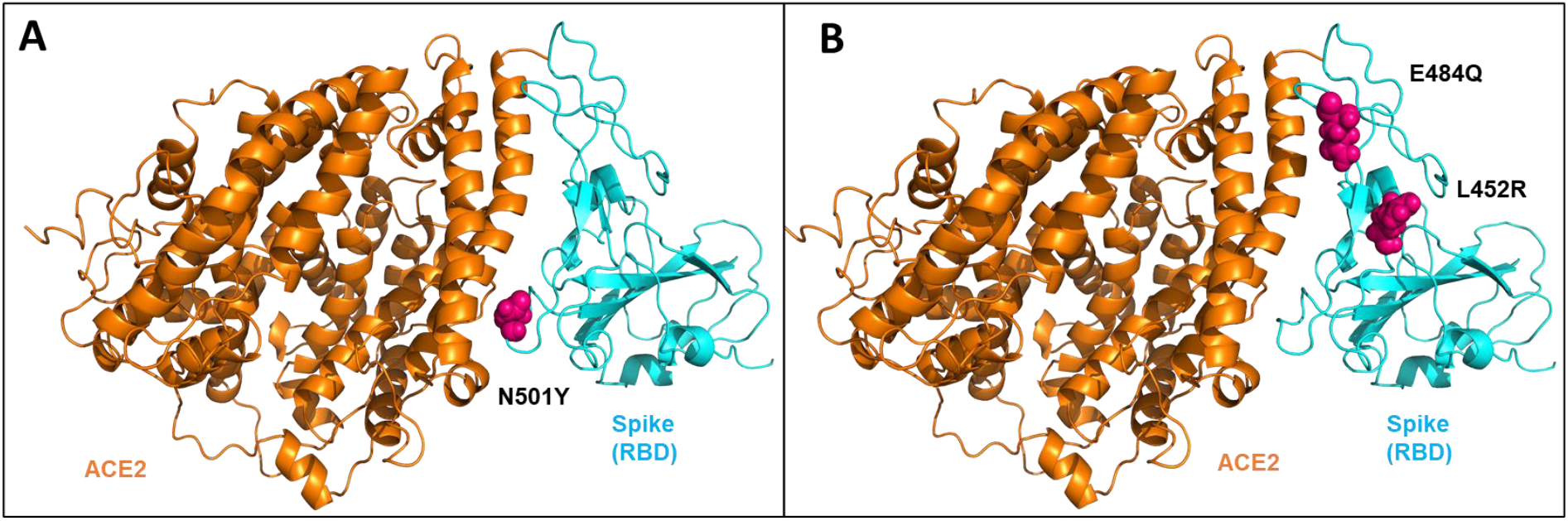
The structure of the Receptor Binding Domain (RBD) of SARS-CoV-2 Spike protein complexed with human Angiotensin Converting Enzyme 2 (hACE2) receptor. (A) The sphere shape residues in hot pink colour show N501Y mutation in Spike protein of SARS-CoV-2. (B) The Double mutation at L452R and E484Q in Spike protein.

In the recent study it was shown that E484 involved in the hydrogen bonding with S2M11 neutralizing antibody and mutation at E484Q/K abrogated the S2M11 mediated neutralization^19^. The B.1.618 (triple mutant), recently detected in the four Indian states (Maharashtra, Delhi, West Bengal and Chhattisgarh), has been characterized by the deletion of TYR145 and HIS146 as well as E484K and D614G mutation in the Spike glycoprotein ^20^ (https://cov-lineages.org/). _Although the sudden spike in COVID-19 cases in India is believed to be because of these two new variants and that might be due to the higher binding affinity towards hACE2 and immune escape ability, there are no biochemical and biophysical studies reported on them yet.

In this study, our aim was to study the thermodynamic effects of the mutations in the RBD region of the Spike glycoprotein interacting with hCAE2 and compare that with the wildtype. We accordingly studied two crucial variants B.1.1.7 and B.1.617, which caused the spike in the COVID19 cases in various countries including India. These two variants possess significant mutations in the RBD domain of the Spike glycoprotein and have a higher infectivity rate. Therefore, to study and compare the dynamics, interactions and binding free energy of Spike protein variants with hACE-2 at the molecular level, we performed the classical Molecular Dynamic (MD) simulations.

## Methods

### MD Simulations

The X-ray crystal structure of SARS-CoV-2, Spike RBD bound with hACE2 was retrieved from Protein Data Bank (PDB) having PDB ID 6M0J. Along with wildtype, two mutants of Spike protein were created, namely, B.1.1.7 (N501Y) and B.1.617 (L452R and E484Q) using the Maestro suite of Schrodinger software ^21^. All the three structures were then pre-processed for missing side chains, deleting waters, the addition of hydrogens, hydrogen bond optimization and restrained minimization using the Protein preparation wizard of Schrodinger software ^21^. The prepared mutated structures were then simulated for 50ns and the last frame structure was taken for further studies. The following protocol was adopted for the MD simulations of all three prepared structures - each system was solvated with the TIP3P water model in an orthorhombic periodic boundary box. To prevent interaction of the protein complex with its own periodic image, the distance between the complex and the box wall was kept 10 Å. The system was then neutralized by the addition of appropriate number of Na+/Cl− ions depending on the protein–ligand complex using OPLS3e forcefield. Then the energy of the prepared systems was minimized by running 100 ps low-temperature (10K) Brownian motion MD simulation (NVT ensemble) to remove steric clashes and move the system away from an unfavourable high-energy conformation. Further, the minimized systems were equilibrated in seven steps in NVT and NPT ensembles using the “relax model system before simulation” option in the Desmond Schrodinger suite ^21^. The equilibrated systems were then subjected to 200 ns unrestrained MD simulations in NPT ensemble with 300 K temperature maintained by Nose–Hoover chain thermostat, constant pressure of 1 atm maintained by Martyna–Tobias–Kelin barostat and an integration time step of 2 fs with a recording interval of 200 ps.

### Analysis of the MD simulation

The Root Mean Square Deviation (RMSD), Root Mean Square Fluctuation (RMSF), number of hydrogen bonding was calculated using the simulation event analysis tool of the Desmond suite integrated into Schrodinger software. Further, the occupancy of the hydrogen bonding between Spike protein and hACE2 was calculated using Visual Molecular dynamics (VMD) software ^22^. The Molecular mechanics generalized born surface area (MM/GBSA) free binding energy between Spike protein and hACE2 was calculated using the prime module of Schrodinger software ^21^. Twenty structures extracted from 50ns to 200ns from each of the trajectories were used for this computation using the following equation:

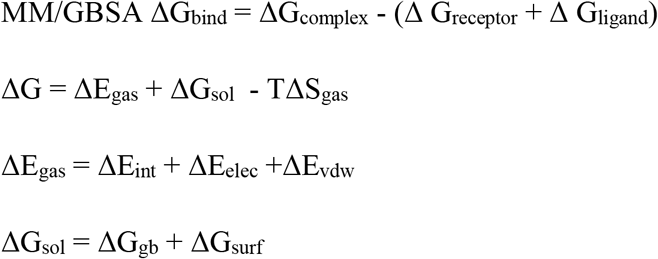

The binding free energy (ΔG_bind_) is dissociated into binding free energy of the complex, receptor and ligand. The gas-phase interaction energy (ΔE_gas_) was calculated as the sum of electrostatic (ΔE_elec_) and van der waal (ΔE_vdw_) interaction energies, while internal energy was neglected. The solvation free energy (ΔG_sol_) contains non-polar (ΔG_surf_) and polar solvation energy (ΔG_gb_), which was calculated by using the VSGB solvation model and OPL3e force field, while the entropy term was neglected by default ^21, 23^.

The energy contribution of the mutated residues was then compared with wildtype residues. The *Prime* module of Schrodinger software was used for the calculation of the energy contribution of the individual residues. The solvent model used here was Surface Generalized Born (SGB), with variable dielectric enabled, the internal dielectric was 1.00 and solvent dielectric was 80.00 ^21^.

The following equation was used for the calculation of prime energy of individual residues:

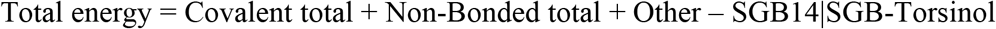

Here, other energy = SGB self, Nonpolar, Hydrogen bond, Packing, self-contact.

## Results and Discussion

### The mutant spike proteins have a better binding affinity with hACE2 in comparison to wildtype

The wildtype (WT) Spike-hACE2 complex, along with the prepared and equilibrated B.1.617 and B.1.1.7 spike variant, were simulated for 200ns. All the three structure complexes were first analysed for investigating the dynamics. In RMSD analysis, we found that all the complexes had a similar deviation around 2.5 Å from the initial structure over the 200ns of simulations; B.1.617_Spike-hACE2 (2.36 ± 0.27 Å), N501Y_Spike-hACE2 (2.62 ±0.67 Å) and WT_Spike-ACE2 (2.82 ± 0.84 Å) as shown in **Figure 2A**. When RMSF of the simulated complexes were analysed, it was found that the residues number ARG355 to PHE400 of spike protein was more flexible, however, no significant difference was found when WT was compared with both the variants. Also, the fluctuation in the mutant residues was not high, although, in the case of B.1.617, it was found that residue VAL445 had more fluctuations than WT and N501Y mutant (**Figure 2B**). The average RMSF for WT was 2.95 ±0.86 Å, for B.1.617, it was 2.66 ± 0.94 Å and for N501Y Spike protein, it was 3.02 ±. 1.09 Å. After analysing the fluctuation and deviations in the structures, the number of hydrogen bond count was calculated between the Spike protein and hACE2 for all three structures. It was found that WT (12.23 ± 2.58) had the highest number of hydrogen bonds, followed by B.1.617 (11.81 ± 2.07) and B.1.1.7 (9.19 ± 1.81), respectively (**Figure 2C**).

**Figure 2.**
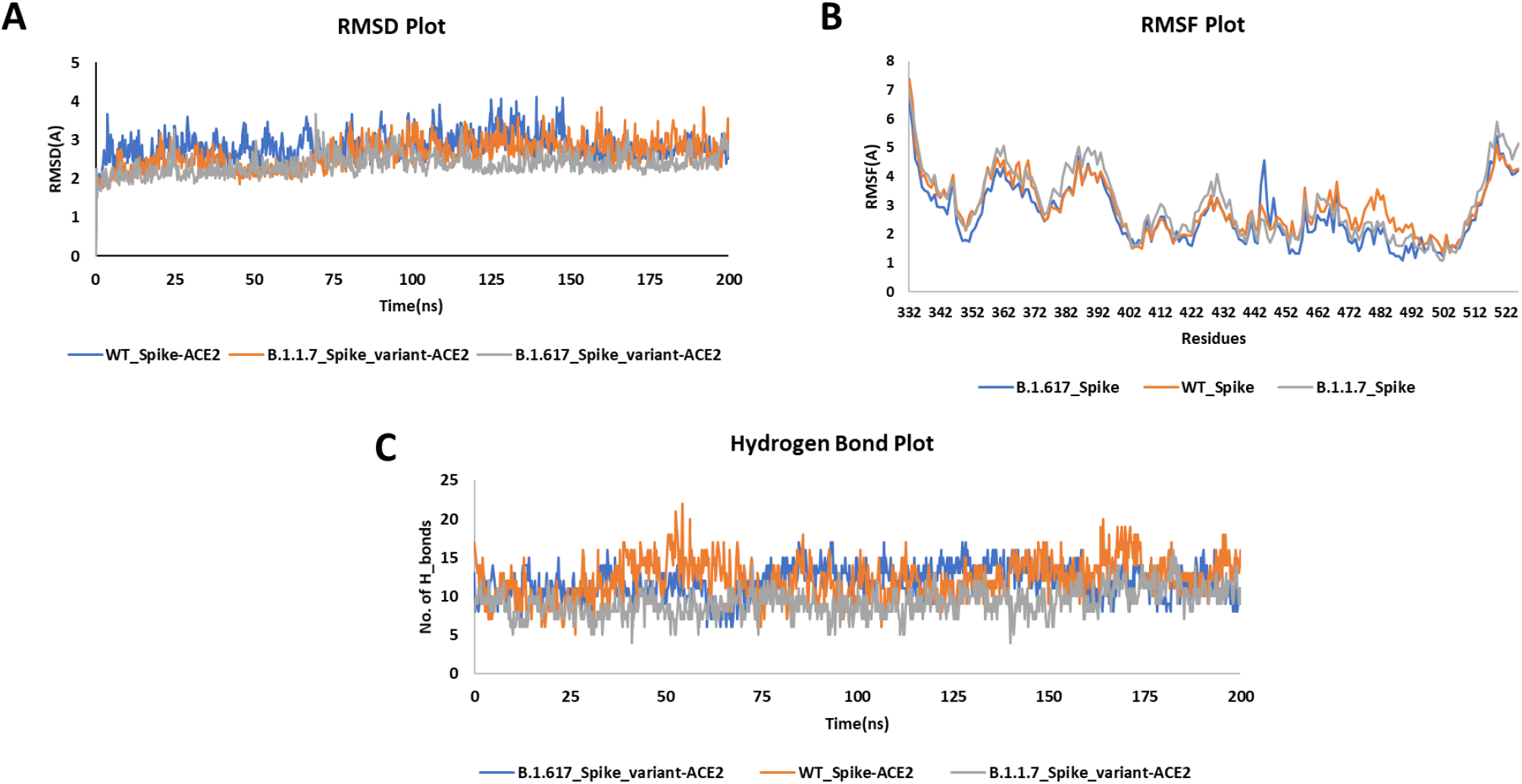
The MD simulation analysis of the three simulated complexes. (A) The RMSD plot showing similar deviation of all the simulated structures. (B) The RMSF plot reveals that residues 350-400 of spike are more flexible, while the mutated residues have lesser fluctuation and are also comparable in all the three structures. (C) The number of hydrogen bond count indicates that WT spike-hACE2 and B.1.617 have similar and higher number hydrogen bonds compared to B.1.1.7 variant.

It was noted that the number of hydrogen bonds was similar in case of WT and B.1.617. We further analysed the significant residues, to find out which of them have greater than 30% of the occupancy of hydrogen bond throughout the simulation. It was found that B.1.1.7 and B.1.617 had more residues interaction than WT. In the case of WT, only three residues (TYR453, THR500 and GLY502) of Spike protein were making hydrogen bond with hACE2 for more than 30% of simulation time. In comparison, in the B.1.1.7 Spike mutant, five residues (LYS417, AlA475, ASN487, THR500 and GLY502) were involved, and in B.1.617, there were six residues (LYS417, TYR449, ASN487, TYR489, THR500 and GLY502) of spike protein that had significant hydrogen bond interactions with hACE2 (**Figure S1**). It was observed that THR500 and GLY502 were the critical residues interacting significantly with hACE2 in all three complexes. None of the mutated residues in B.1.617 (L452R, E484Q) or N501Y were found to be interacting significantly with hACE2. Hence, it was essential to investigate if these mutated residues of Spike protein had interaction with any other Spike residues or any interaction with hACE2 for any fraction of time. To analyse the changes in the interaction due to mutation, we extracted the three structures at the 50ns interval from all the three simulated complexes. It was found that in the case of B.1.617, in the 50^th^ ns frame, neither ARG452 nor GLN484 was involved in any polar contact with other residues, while in 100^th^ ns and 150^th^ ns frame, it was found that GLN484 was making hydrogen bond contact with SER349 and ASN450 of Spike protein itself, while in WT Spike protein, GLU484 was making hydrogen bond only with SER439. So, it was observed that there was an increase in intra-chain interaction of Spike protein due to mutation of E484Q. Similarly, when B.1.1.7 variant was compared with WT, it was found that due to ASN to TYR mutation at 501^th^ residue, there was increase in the hydrogen bonding with LYS353 of hACE2 (**Figure 3**). The increase in the intra-chain interaction in case of B.1.617 indicated that it may interfere in the human antibodies interaction with Spike protein, which may lead to higher infectivity rate. While, in case of B.1.1.7 the increase in hydrogen bond contact with hACE2 indicated the higher binding affinity of this mutant with hACE2 in comparison to WT.

**Figure 3.**
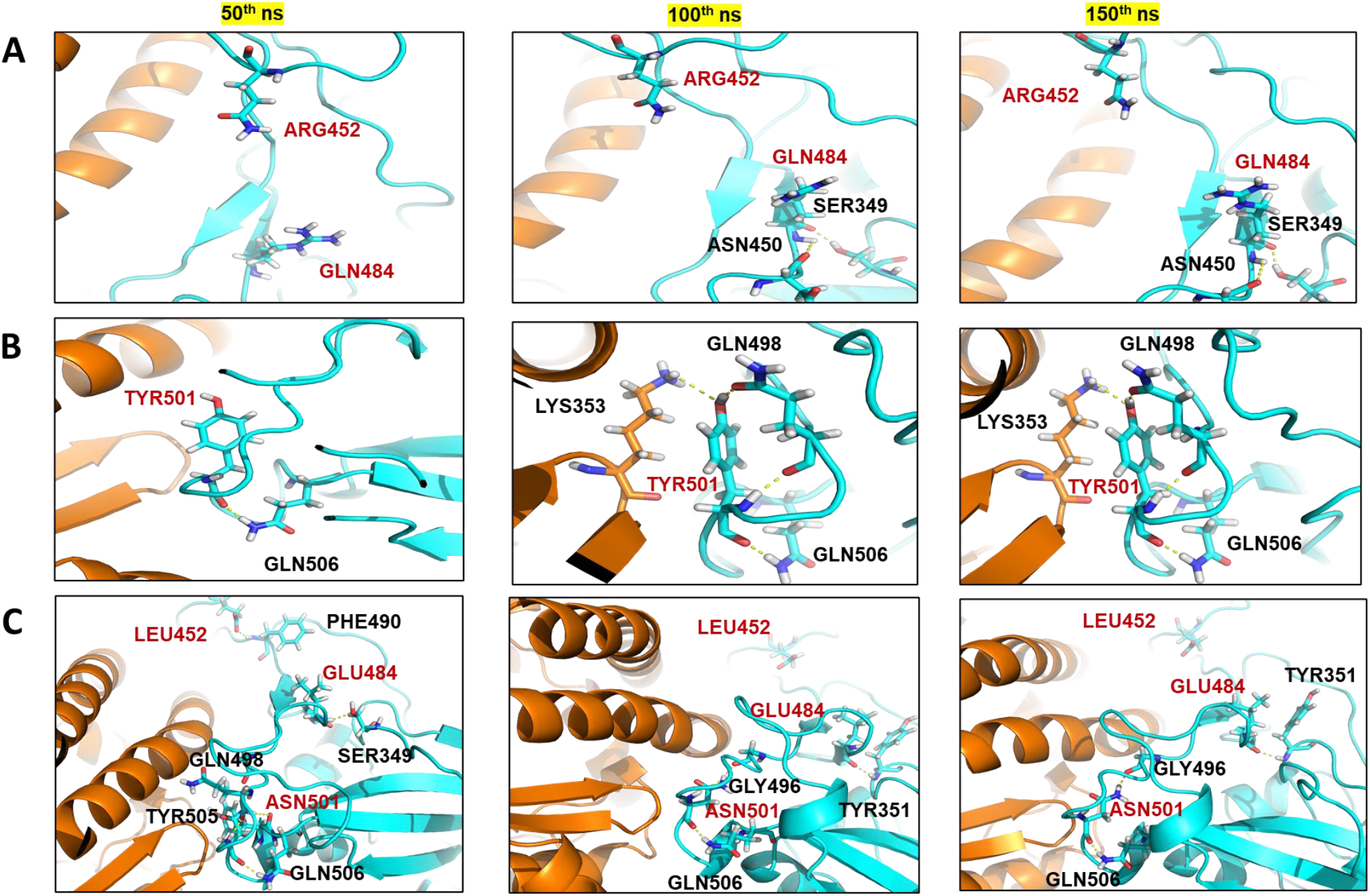
The comparison of the interaction of the mutated residues and wild-type residues in the three structures extracted at 50ns span from the simulated trajectories. (A) B.1.617 (L452R and E484Q) spike variant making polar interactions at 50^th^ ns, 100^th^ ns and 150^th^ ns, respectively. (B) The interaction of Tyrosine at 501st position of mutant Spike protein (C), The interaction of wild type residues at 50^th^, 100^th^ and 150^th^ ns of the simulation.

The MM/GBSA binding free energy has been earlier reported in literature to correlate with the binding affinity between the complexes ^21, 24^. Therefore, to assess the affinity of Spike protein towards hACE2, we calculated the MM/GBSA binding free energy by extracting twenty structures in equal span from 50^th^ to 200ns of the simulated trajectories. The MM/GBSA binding free energy showed that mutants Spike proteins had higher binding affinity with hACE2 than WT. It was found that B.1.1.7 (103.35 ± 16.31 kcal/mol) and B.1.617 (101.90 ± 18.40 kcal/mol) Spike proteins had a similar binding affinity with hACE2 in comparison to WT (−96.87 ± 14.57 kcal/mol) (**Figure S1**).

Further, to calculate the energy contribution of individual mutated residues, prime energy was calculated for the twenty extracted structures, which showed that B.1.617 (L452R, E484Q) had a more stabilizing effect on Spike protein comparison to WT. The average energy contribution of ARG (−50.90 ± 3.99 kcal/mol) in comparison to LEU (−22.15 ± 2.60 kcal/mol) at 452^nd^ position of Spike protein was found to be high. Similarly, the energy contribution of GLN (−56.03 ±2.31 kcal/mol) in comparison to GLU (−47.12 ± 2.25 kcal/mol) at 484^th^ position of Spike protein was quite higher. However, in the case of the B.1.1.7 mutant, it was noticed that ASN (−64.31 ± 3.59 kcal/mol) at 501^st^ position was more energetically favourable than TYR (−31.55 ± 3.26 kcal/mol) (**Table1**). Although these binding energy calculations are theoretical and cannot be taken as absolute values, however, they are typically used for the comparison of the binding affinity of the complexes with respect to each other. The interactions and binding energy calculations showed that in B.1.617, there is an increase of energy due to mutation as well as change in intra-chain interaction, which may lead to stabilization and change in the orientation of Spike protein.

**Table 1.** The residue wise energy contribution of the mutated residues compared with the wildtype for the twenty structures extracted from 50 to 200 ns of the simulation for all the three complexes.

| **B.1.617_Spike(kcal/mol)** |  | **WT_Spike protein (kcal/mol)** |  |  | **N501Y_Spike (kcal/mol)** |
| --- | --- | --- | --- | --- | --- |
| **R452** | **Q484** | **L452** | **E484** | **N501** | **Y501** |
| -47.16 | -51.43 | -23.45 | -47.06 | -65.7 | -31.72 |
| -50.69 | -57.57 | -23.04 | -48.53 | -67.57 | -27.6 |
| -42.13 | -56.15 | -23.89 | -45.52 | -66.44 | -32.73 |
| -52.43 | -56.51 | -20.05 | -48.94 | -63.37 | -31.96 |
| -54.38 | -58.47 | -23.68 | -50.2 | -61.83 | -26.71 |
| -54.05 | -56.86 | -25.88 | -47.91 | -67.16 | -29.47 |
| -56.11 | -53.98 | -21.65 | -47.86 | -60.13 | -37.94 |
| -47.14 | -56.4 | -22.44 | -44.1 | -65.41 | -33.03 |
| -56.47 | -57.21 | -22.87 | -43.55 | -61.79 | -31.87 |
| -54.79 | -53.25 | -22.05 | -48.18 | -61.17 | -33.2 |
| -50.07 | -51.51 | -20.9 | -46.46 | -60.9 | -33.69 |
| -45.38 | -53.04 | -17.67 | -45.08 | -60.25 | -25.84 |
| -49.22 | -59.17 | -25.26 | -49.82 | -66.88 | -34.98 |
| -51.75 | -58.8 | -16.19 | -47.49 | -66.96 | -29.93 |
| -45.95 | -55.64 | -18.58 | -47.46 | -57.8 | -34.11 |
| -48.85 | -56.21 | -20.55 | -49.06 | -61.5 | -34.51 |
| -48.93 | -56.69 | -24.27 | -46.24 | -64.05 | -31.49 |
| -54.01 | -56.35 | -23.78 | -50.55 | -68 | -32.64 |
| -55.01 | -59.25 | -21.35 | -41.95 | -67.16 | -32.55 |
| -53.5 | -56.27 | -25.55 | -46.55 | -72.31 | -25.06 |
| **-50.90± 3.99** | **-56.03± 2.31** | **-22.15± 2.60** | **-47.12± 2.25** | **-64.31± 3.59** | **-31.55± 3.26** |

Recently it was also reported about the abrogation of interactions with neutralizing antibody S2M11 due to E484K mutation ^19^. Overall, stabilization of spike protein and increase in intrachain interactions and chance of immune scape could be helping the B.1.618 variant to be more transmissible. While in the B.1.1.7 Spike mutant, an increase in the hydrogen-bond interaction and binding affinity with hACE2 could be the reason for more transmissivity of this mutant.

## Conclusion

In this study, we performed the MD simulations and have compared the binding energy, interactions, and change in dynamics of three protein complexes, namely WT, B.1.617 (L452R, E484Q) and N501Y mutant. We showed that mutants have a higher number of significant hydrogen bond interactions with hACE2, and the binding free energy of the mutants is also higher in comparison to WT. In B.1.617, the mutations were energetically favourable as was also observed in terms of intra-chain residue interactions. In B.1.1.7, the mutation leads to an extra interaction with hACE2 (LYS353) in comparison to WT. The detailed molecular level interaction dynamics of Spike-hACE2 interactions and the predicted increased structural stability of its spike protein and hACE2 affinity can be possibly linked to higher transmissibility of B.1.617 and B.1.1.7 variants of SARS-CoV-2

## Notes

### Competing Interest Statement

The authors have declared no competing interest.

