## Supplementary Figure1 for "Possible link between higher transmissibility of B.1.617 and B.1.1.7 variants of SARS-CoV-2 and increased structural stability of its spike protein and hACE2 affinity"

**Supplementary Figures**

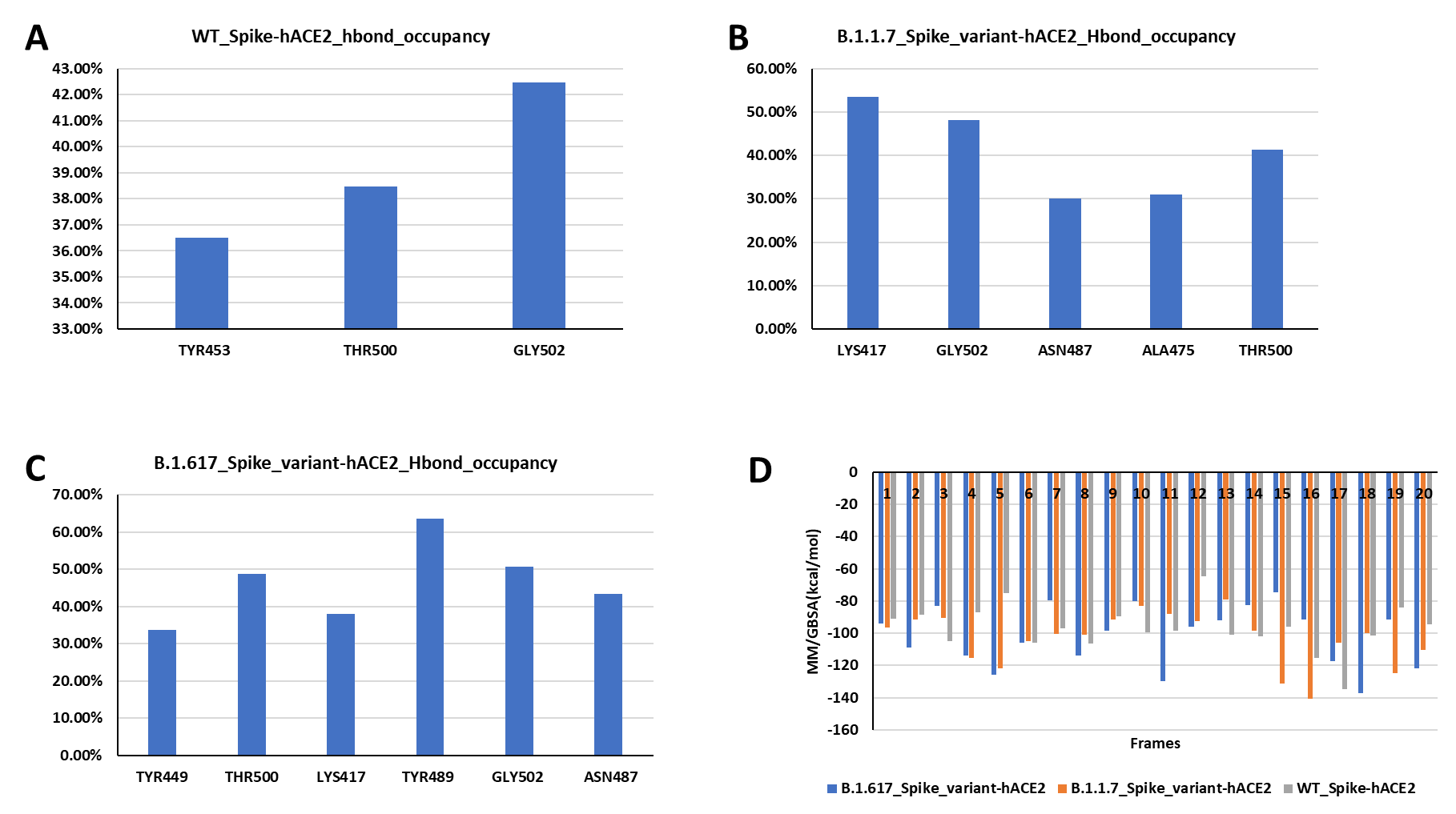

**Figure S1.** (A) The hydrogen bond occupancy in WT spike-hACE2 complex, (B) B.1.1.7 variant and (C) B.1.617 throughout the 200ns of MD simulations. (D) MM/GBSA free binding energy for the twenty structures extracted at equal span from 50 to 200ns of simulated trajectories for all the three complexes.
